# Targeting Spike Glycans to Inhibit SARS-CoV2 Viral Entry

**DOI:** 10.1101/2022.12.22.521642

**Authors:** Alex J. Guseman, Linda J. Rennick, Sham Nambulli, Chandra N. Roy, David R. Martinez, Darian T. Yang, Fatema Bhinderwhala, Sandra Vergara, Ralph S. Baric, Zandrea Ambrose, W. Paul Duprex, Angela M. Gronenborn

## Abstract

SARS-CoV-2 Spike harbors glycans which function as ligands for lectins. Therefore, it should be possible to exploit lectins to target SARS-CoV-2 and inhibit cellular entry by binding glycans on the Spike protein. *Burkholderia oklahomensis* agglutinin (BOA) is an antiviral lectin that interacts with viral glycoproteins via N-linked high mannose glycans. Here, we show that BOA binds to the Spike protein and is a potent inhibitor of SARS-CoV-2 viral entry at nanomolar concentrations. Using a variety of biophysical tools, we demonstrate that the interaction is avidity driven and that BOA crosslinks the Spike protein into soluble aggregates. Furthermore, using virus neutralization assays, we demonstrate that BOA effectively inhibits all tested variants of concern as well as SARS-CoV 2003, establishing that glycan-targeting molecules have the potential to be pan-coronavirus inhibitors.

## Introduction

The pandemic of coronavirus disease 2019 (COVID-19) has claimed over 6 million lives to date (*1*). In the race for pharmacological interventions, the Spike protein on SARS-CoV-2, the causative agent of COVID-19, has become the predominant target for vaccine development and early monoclonal antibody therapeutics (*2–4*). The Spike protein forms a trimer, approximately 25 of which protrude from the outer surface of the virion in a semi-stabilized prefusion state (*5, 6*). Upon engagement with the cellular receptor, angiotensin converting enzyme-2 (ACE2), Spike undergoes a substantial conformational change, triggering fusion between the virus and host cell membranes. This process is initiated by and requires the transition of the receptor binding domain (RBD) from a buried “down” state, to an exposed, active “up” state. Although antibodies are effective weapons in the fight against COVID-19, new variants of SARS-CoV-2 are rapidly evolving in the race between virus and host, rendering many antibodies ineffective (*7*).

Like other viral envelope proteins, Spike is heavily glycosylated, and this glycan shield cloaks the protein, effectively hiding 44% of its solvent accessible surface area from recognition by the host immune system (*8–11*). Spike sugars are predominately N-linked and have been mapped to 22 different asparagine (Asn) residues (*8, 11, 12*). They range in glycotype from high mannose to complex glycans, varying for each glycosylation site (*8*). Although most glycans appear to hide the protein surface, there is growing evidence that some possess additional structural roles (*13, 14*). In particular, virions produced from CRISPR engineered cells with perturbed glycan processing machinery and altered glycosylation patterns exhibit reduced infectivity (*15*). Furthermore, an all atom molecular dynamics simulation of glycosylated Spike suggested that the N165 and N234 glycans regulate the up/down RBD conformational switch (*13*), while the N343 glycan acts as a gate to permit the up/down placement of the RBD (*14*). Subsequent biophysical and structural studies, in which the glycosylation capacity of specific residues was ablated by selective mutation, confirmed a pivotal role for the N165 and N234 glycans in RBD positioning and ACE2 binding (*16*). Therefore, targeting and exploiting glycans on Spike constitutes an attractive strategy for developing SARS-CoV-2 inhibitors.

Glycans are recognized and bound by lectins, a class of proteins that is present in all domains of life, ranging from bacteria and fungi to plants and animals (*17*). Importantly, most lectins possess exquisite configurational and conformational specificity towards particular glycans. Functionally, lectins play key roles in a wide variety of biological processes, including host pathogen interactions, cell-cell signaling, and immune modulation (*18*). Among the antiviral lectins, activities against a wide range of viruses have been reported, including HIV, Ebola virus, SARS-CoV 2003, and MERS-CoV (*17, 19, 20*). As potential therapeutics, however, their uses have not been fully realized (*17, 19–22*)

Structurally, lectins adopt a diverse set of architectures (*17, 22–25*). They can bind glycans mono- or multivalently at single or multiple sites. The anti-HIV activities of lectins are associated with binding to the high-mannose glycans on gp120, the major HIV envelope glycoprotein that is critical for entry of the virus into the host cell. Among the HIV-inactivating lectins, the *Oscillatoria agardhii* agglutinin (OAA) family exhibits an unprecedented beta barrel topology, in single domain or multi-domain arrangements (*22*). Each beta barrel contains two glycan binding sites, targeting the core 3α,6α-mannopentaose of N-linked high mannose glycans on the HIV envelope protein and inhibiting HIV infection at nanomolar concentration (*21, 22, 24, 26*). For example, the OAA family member *Burkholderia oklahomensis* agglutinin (BOA) has a dual beta barrel topology and an effective EC_50_ concentration against HIV of 12 ±2 nM (*24*). Since binding of 3α,6α-mannopentaose by individual BOA binding sites occurs with K_d_ values between 20-60 μM, as measured by NMR, the high apparent inhibitory activity of BOA is avidity driven, through multiple interactions with gp120 (*24*).

Here, we demonstrate that BOA interacts with the SARS-CoV-2 Spike protein and inhibits cellular entry of SARS-CoV-2 at concentrations less than 50 nM.

## Results

### Lectins as inhibitors of viral entry

Three different lectins, griffithsin (GRFT), OAA, and BOA, were evaluated for their ability to inhibit infection of Vero E6 cells by the Munich SARS-CoV-2 authentic virus isolate. All three lectins have been reported to inhibit HIV-1 infection, although GRFT and OAA/BOA differ significantly in structure and glycan binding specificities (*22, 24, 25*). No inhibition of SARS-CoV-2 infectivity was observed for GRFT (at 80 nM), partial inhibition was seen for OAA (at 75 nM), and complete inhibition was achieved using BOA (at 115 nM) (Figure S1).

Since BOA was the most potent lectin tested, we evaluated its activity in three different virus neutralization systems. First, using plaque assays, we found that BOA blocked the infectivity of the SARS-CoV-2 Munich isolate in Vero E6 cells with an EC_50_ value of 9 ±2 nM (Figure 1C). Second, using replication-defective, luciferase-expressing HIV-1 pseudotyped with Spike from the Wuhan-Hu-1 isolate containing the D614G substitution, BOA reduced infection of Vero E6 cells, as measured by luciferase activity, with an EC_50_ value of 1.6 ±0.3 nM (Figure 1D). Note, the D614G substitution increases the Spike protein density on virions resulting in improved transduction (*27–29*). Finally, infection of HEK293F cells expressing human ACE2 (hACE2) with replication-incompetent GP-expressing HIV-1 pseudotyped with Wuhan-Hu-1 D614G Spike, generated an EC_50_ value of 48 ±16 nM (Figure S2). All three assays demonstrate that BOA inhibits SARS-CoV-2 cellular entry at nanomolar concentrations.

**Figure 1:**
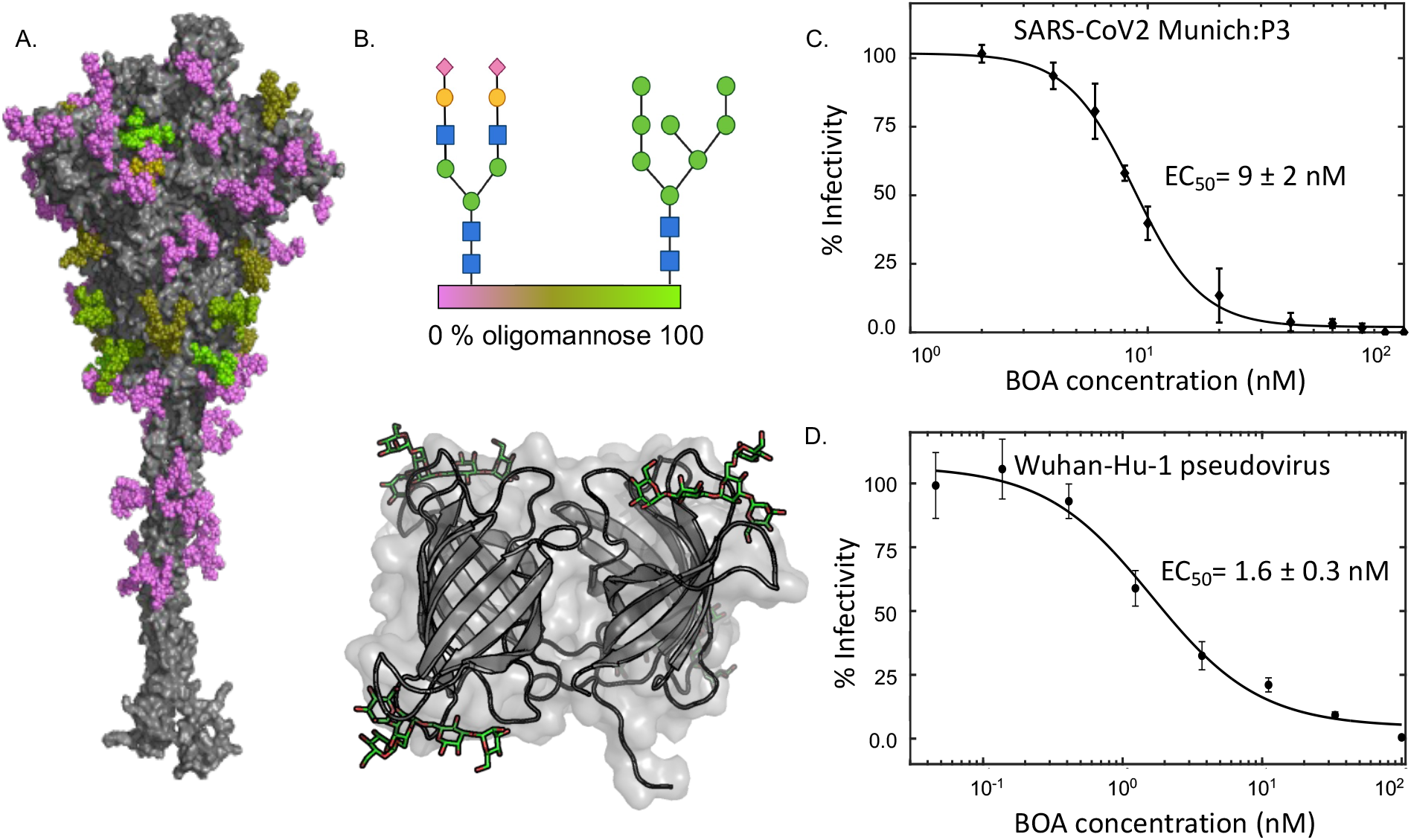
Inhibition of SARS-CoV-2 infection by BOA. A. Surface representation of the SARS-CoV-2 Spike protein (gray) with glycans colored according to glycotype (>90% oligomannose occupancy (lime), 50-75% oligomannose occupancy (olive), or complex (violet)). B. Schematic representation of a complex glycan and high mannose glycan (top) and the BOA-mannopentaose structure (bottom). Each sugar is denoted by shape and color (green circles, mannose; blue squares, N-acteyl glucosamine; yellow circles, galactose; purple diamonds, N-acetyl-neuraminic acid). The structure of BOA is shown in ribbon representation and 3α,6α-mannopentaose in green stick representation (PDB ID: 4GK9). C. Concentration-dependent inhibition of the SARS-CoV-2 Munich isolate by BOA, as measured by a plaque reduction assay and following normalization, yielding an EC_50_ value of 9 ±2 nM. D. Concentration-dependent inhibition of HIV-1 pseudotyped with Wuhan-Hu-1 Spike protein by BOA, as measured by using a luciferase reporter, yielding an EC_50_ value of 1.6 ±0.3 nM.

### Biophysical Characterization of Lectin Spike interactions

To explore whether BOA and OAA binding to Spike was responsible for inhibition of infection, *in vitro* binding was evaluated using purified Spike, OAA, and BOA proteins. Mixing equimolar amounts of lectin and Spike proteins, at micromolar concentrations, resulted in co-precipitation (Figure S3). Size exclusion chromatography (SEC) generated data consistent with this observation. Specifically, Spike protein alone eluted at a volume of 13 mL (Figure 2A and B), consistent with its molecular mass of ~500 kDa and previous reports (*30, 31*). In the presence of OAA or BOA, each at either 5:1 or 10:1 molar excess, additional elution peaks were observed, at ~8 mL (Figure 2A and B), the exclusion limit of the column, suggesting that large molecular mass complexes are formed. Visualization of the material in the peak fractions by negative stain electron microscopy revealed large amorphous aggregates (Figure S4). Building on these observations, we characterized the concentration dependence of OAA- and BOA-induced aggregation of Spike protein by dynamic light scattering (DLS). At concentrations as low as 60 nM BOA or 125 nM OAA, a significant increase in the hydrodynamic radius of the Spike protein complex was observed. This increase in hydrodynamic radius continued with increasing lectin concentration until the detector limit was reached (Figure 2C). Finally, we used SeTau-405 fluorescently labeled BOA in fluorescence titration experiments with Spike. In agreement with the above results, saturable binding was observed with a half-saturation point between 30-60 nM (Figure 2D). This value is the same order of magnitude as the measured inhibitory concentrations BOA in infectivity assays.

**Figure (2):**
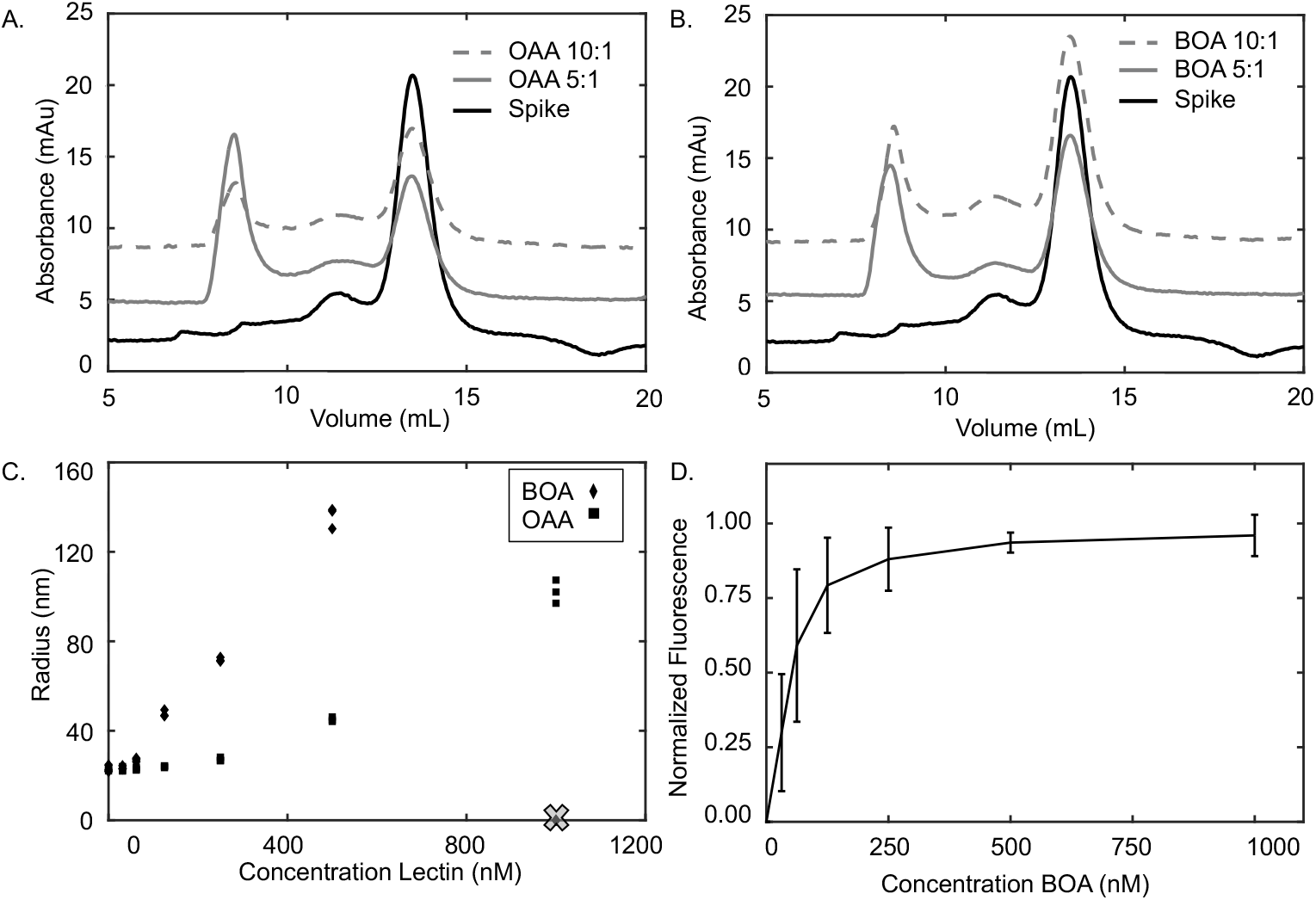
OAA and BOA interact with and aggregate Spike protein *in vitro* at nM concentrations. A and B. Size exclusion chromatography of Spike protein without or with OAA (A) or BOA (B) at 5:1 (gray) and 10:1 (gray dotted) molar excess of lectin. C. Hydrodynamic radius of particles monitored by DLS of Spike protein in the presence of increasing concentrations of OAA or BOA. D. Binding isotherm of BOA binding to Spike protein extracted from fluorescence measurements. Normalized fluorescence is plotted as a function of BOA concentrations.

### Mechanism of Inhibition

The high mannose glycan on N234 of the Spike protein has been implicated in regulating the position of the RBD (*13, 16*). We therefore considered the possibility that BOA binding to this particular glycan plays a role in its inhibition of viral entry. To test this idea, we generated pseudoviruses expressing Spike protein variants N234A and as a control N165A. While the N165A variant infected Vero E6 cells, the N234A variant was noninfectious (Figure S5), confirming that the identity of N234 and/or the glycan at this position is important for viral entry. As N165 is a complex glycan and thus not targeted by BOA, deletion of this glycan by the N165A mutation had minimal impact on BOA’s ability to inhibit viral entry, resulting in an EC_50_ value of 10 ± 2 nM (Figure S5D). Given that the N234A pseudovirus was not infectious, N234A Spike and BOA interactions were assessed *in vitro* using size exclusion chromatography. Similar to what was observed with WT Spike protein, BOA and N234A Spike eluted at ~8 mL, the exclusion limit of the column, consistent with the formation of soluble aggregates. To corroborate our experimental observations, we computationally evaluated the solvent accessibility of the glycans on the Spike protein structure using different size probes, ranging in radius from 1.4 Å, comparable to a water molecule, to 20 Å, comparable to the size of BOA (Figure S6). The data indicate that the N234 glycan is buried in the WT Spike structure and that BOA interacts with the N234A Spike variant through multiple glycans. It is likely that N234 glycan is inaccessible to BOA and does not play an important role in BOA-mediated inhibition.

The role of multivalency in the interaction between BOA and Spike was tested by removing three of the four glycan binding sites on BOA. Key arginine, glutamic acid, and tryptophan residues were substituted by alanine. The resulting BOAd3WRE1 variant (W18A:R101A:E64A:W218A:R170AE197A:W152A:R261A:E236A) is a monovalent version of BOA (Figure 3A) that retains the wild-type (WT) BOA fold and is capable of sugar binding, as evidenced by NMR (Figure S7). DLS measurements revealed that the BOAd3WRE1 variant fails to aggregate Spike at nanomolar concentrations (Figure 3B). Similarly, SEC of Spike in the presence of BOAd3WRE1 shows that the elution volume of the complex is only slightly changed from that of Spike alone, indicating that no large soluble aggregates are formed (Figure 3C). In agreement with the above *in vitro* data, BOAd3WRE1 failed to interfere with infectivity of the authentic SARS-CoV-2 or pseudovirus (Figure 3D and E), demonstrating the importance of multivalency in the interaction between OAA and Spike for antiviral activity.

**Figure 3.**
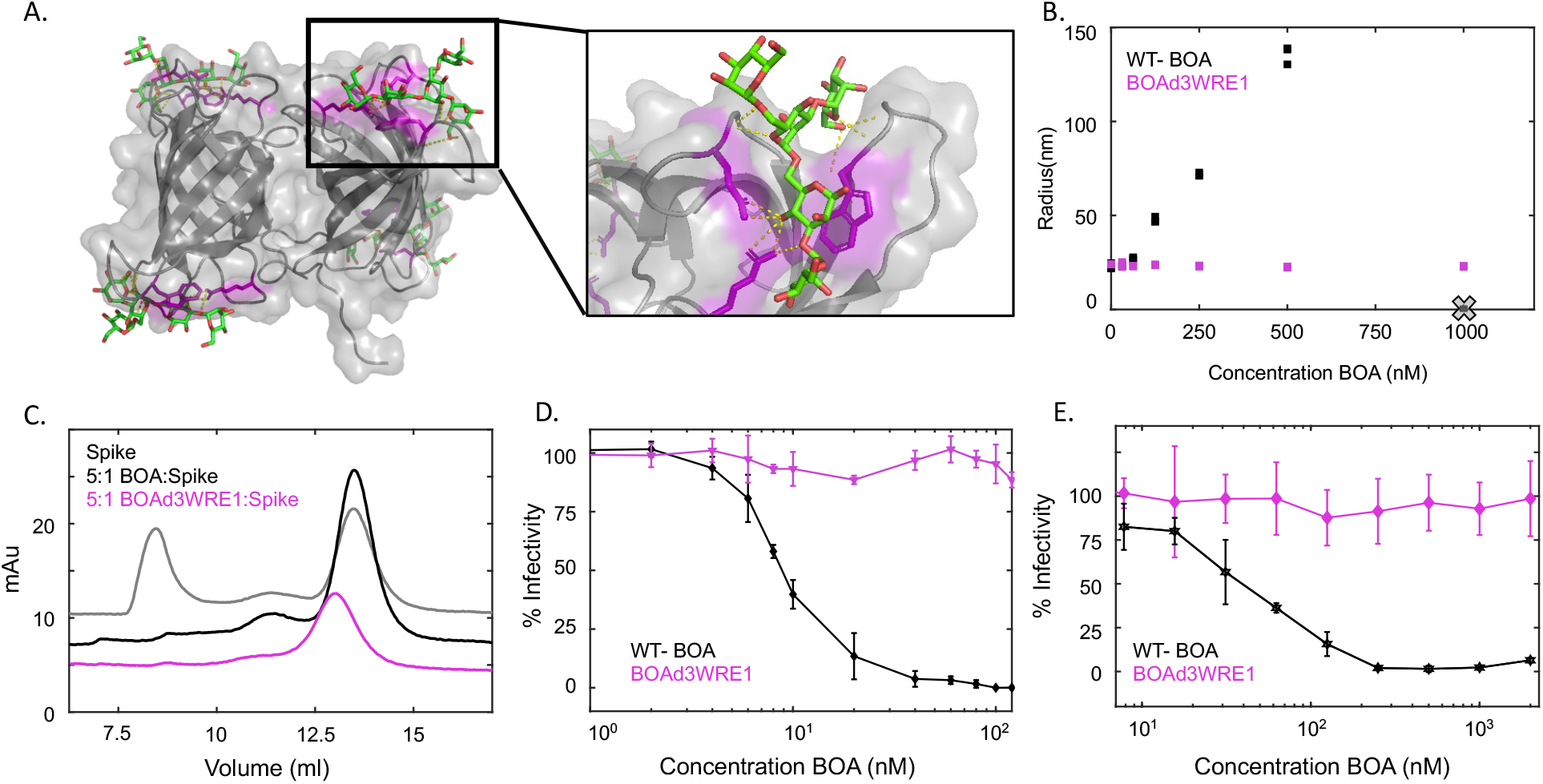
Glycan binding sites in BOA and their importance for avidity-mediated interactions with the Spike protein. A. X-ray structure of the BOA-mannopentaose complex in transparent surface and embedded ribbon representation (PDBID 4GK9). Key Arg, Glu and Trp residues that are important for glycan recognition in each binding site are highlighted in purple stick representation with the glycan shown in green stick representation. B. Comparison of Spike protein aggregation in the presence of increasing concentrations of WT BOA or BOAd3WRE1, which lacks three of its four glycan binding sites. No aggregation was observed for the variant with a single binding site. C. SEC of Spike protein alone (black) with WT-BOA (gray trace) or with BOAd3WRE1 (magenta trace) at a 5:1 lectin:Spike protein ratio.. D. and E. Infectivity assays with WT BOA and BOAd3WRE and Vero E6 cells with the SARS-CoV-2 Munich isolate (D) or of HEK293F cells expressing hACE2 and infection by HIV-1 pseudotyped with D614G Spike (E).

### Effectiveness Against Variants of Concern

As new variants of SARS-CoV-2 emerge, mutations that facilitate transmission and/or immune escape are classified by the World Health Organization (WHO) as variants of concern (*7, 33–37*). (Figure 4A). For all variants of concern, decreased recognition by several antibodies has been reported and symptomatic infections in fully vaccinated individuals with these variants have been observed (*7, 36–38*). Numerous mutations in the virus, including to the Spike gene, have been mapped, especially for regions in the protein under selective pressure, such as position 484 in the RBD or position 681 in the furin cleavage site of the Spike protein (*39, 40*). In contrast, all 22 sites of Asn-linked glycans have remained invariant since the original SARS-CoV-2 isolates in 2019 (Figure 4B), suggesting that Spike retains its original glycan shield. Therefore, BOA should be able to interact with these sugars and retain its ability to inhibit cellular entry of these viral variants. This was tested for the Munich isolate, Delta, and Omicron variants, and only negligible changes in EC_50_ were observed (Figure 4C and Table S1). BOA-mediated inhibition was also tested using replication-defective lentiviruses that were pseudotyped with the Spike protein from several variants of concern. Our data show that BOA inhibits Alpha, Beta, Gamma,Delta, and Omicron variants of concern with EC_50_ values similar to that of the ancestral virus (Figure 4D and Table S1).

**Figure 4:**
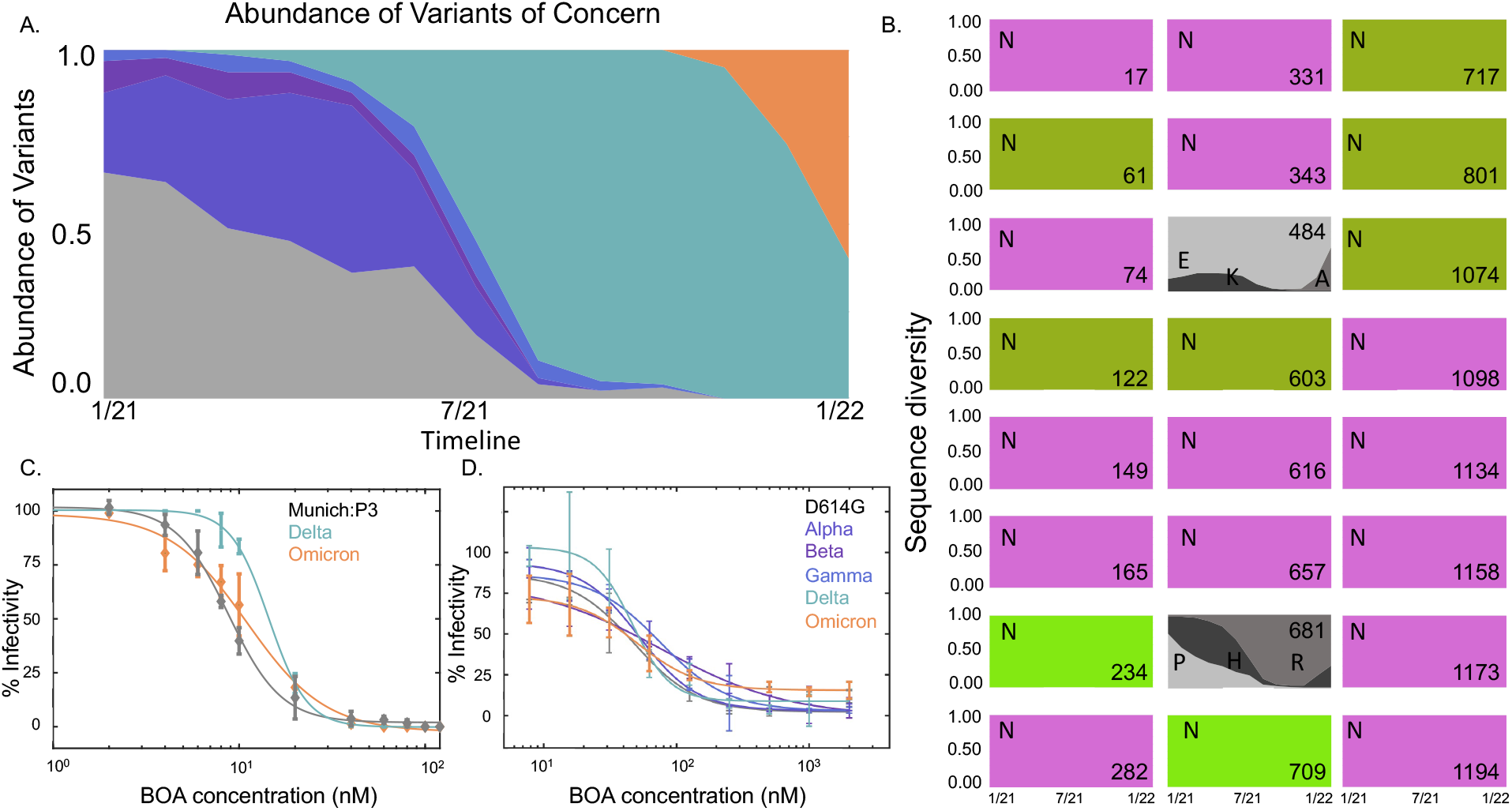
Assessment of BOA activity against emerging variants of concern. A. World-wide prevalence of ancestral SARS-CoV-2 (grey) and Alpha (indigo), Beta (purple), Gamma (blue), Delta (teal), and Omicron (orange) variants of concern in 2021 (Data from Nextstrain database (*32*)). B. Sequence conservation and diversity across variants at positions that possess glycosylated Asn residues. The boxes are colored by glycotype, >90% oligomannose occupancy (lime), 50-75% oligomannose occupancy (olive), complex (violet), and positions are shown by numbers in the bottom right corner. In addition to positions of Asn residues, amino acids at positions 484 in the RBD and 681 in the furin cleavage site are shown in grey boxes with amino acids that changed in variants of concern labeled in one letter code. C and D: Infectivity data for several variants of concern using both authentic SARS-CoV-2 (C; Delta and Omicron) and pseudotyped viruses (D; Alpha, Beta, Gamma, Delta, and Omicron).

### BOA is a potential Pan-CoV inhibitor

Since SARS-CoV-2 and SARS-CoV 2003 are closely related coronavirus with similar Spike protein structures (Figure 5A), exhibiting 70% amino acid sequence identity, we evaluated whether BOA can also inhibit SARS-CoV 2003. Importantly, both Spike proteins possess 22 sites of N-linked glycosylation, with 28% and 32% oligomannose content found in the Spike proteins of SARS-CoV-2 and SARS-CoV 2003, respectively (*8, 9*). Gratifyingly, BOA inhibits infection by SAR-CoV 2003 with an EC_50_ value of 26 ±0.3 nM (Figure 5B), the same order of magnitude as seen for the SARS-CoV-2 variants (cf. Figures 1C, D and 4C, D).

**Figure 5:**
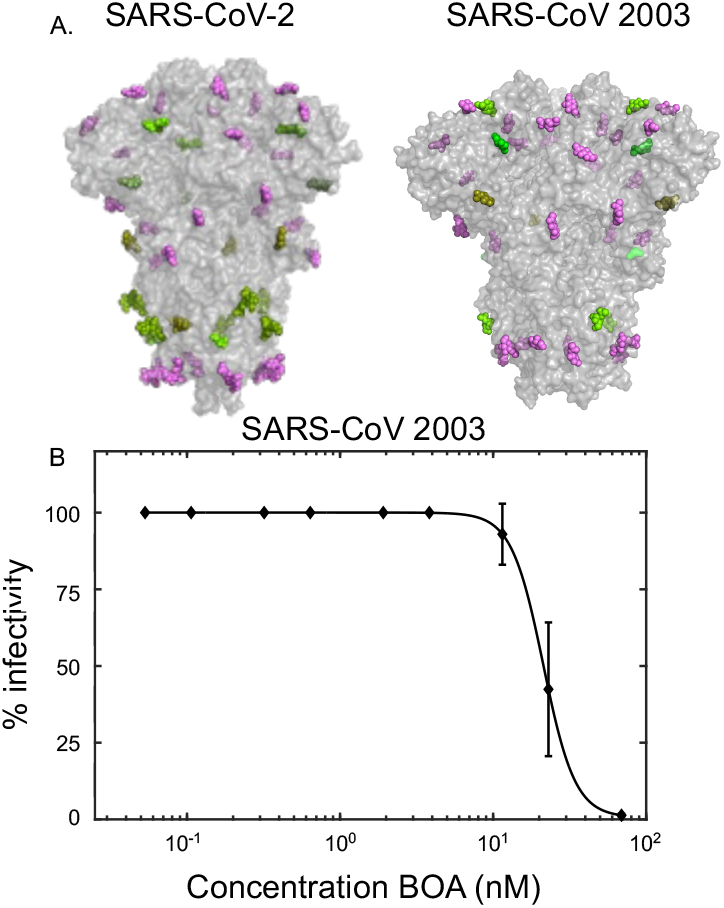
Comparison of SARS-CoV-2 and SARS-CoV 2003 Spike proteins and BOA inhibition of SARS-CoV 2003 A. Surface representation of the X-ray structures of SARS-CoV-2 (PDBID: 6VXX) and SARS-CoV 2003 (PGB ID: 5X58) Spike proteins. Amino acids are shown in gray and sugars are colored by glycotype: >90% oligomannose occupancy (lime), 50-75% oligomannose occupancy (olive), complex (violet). B. Inhibition assay of SARS-CoV 2003 Urbani isolate modified with a luciferase reporter reveals an EC_50_ of 26 ±0.3 nM.

## Discussion

The ability of three lectins (BOA, OAA, and GRFT) to interfere with SARS-CoV-2 infectivity was evaluated employing different assays. While GRFT exhibited no activity in all assays, both BOA and OAA possess potent inhibitory activity. BOA and OAA belong to the same structural family and bind to identical sugar substructures of high mannose glycans (*22, 24*), whereas GRFT is structurally very different and binds to the terminal sugars in the D2 and D3 arms of high mannose glycans (*25*). Unfortunately, conflicting reports on GRFT’s activity towards SARS-CoV-2 infection exist, describing very different EC_50_ values of 6 nM (*41*) or 300 nM (*42*). Since we did not observe any inhibition at 80 nM GRFT concentration, our results are in line with the data reported by the Jiang lab (26). Whether the observed disparities are due to different cellular glycan processing, variations in viral isolates, or specific details in the assays remains to be evaluated.

The key structural difference between OAA and BOA lies in the number of beta barrel domains and the number of glycan binding sites: BOA contains two domains and therefore four glycan binding sites, while OAA has a single beta-barrel with two binding sites. Since anti-viral activity appears to be related to glycan binding, BOA, with twice the glycan binding sites compared to OAA, is a more potent inhibitor. Likewise, removing three of the four sites on BOA obliterates activity, confirming that multi-site interactions are essential. Considering our biophysical assays that report the formation of soluble aggregates at nanomolar concentrations, the multi-site interaction between BOA and the Spike protein most likely accounts for the anti-viral activity of the lectin.

Since one or more glycans on the Spike protein are likely targeted by BOA and given the reported importance of the N234 glycan for the conformation of the Spike RBD that engages the cellular receptor ACE2, we aimed to test whether the N234A Spike glycan deletion variant could be inhibited by BOA. However, because pseudovirus carrying the N234A Spike modification was noninfectious, a specific role for the glycan in BOA-mediated viral inhibition could not be ascertained. Instead, the inability of N234A virus to infect cells suggests that N234 and/or its glycan is essential for viral entry. The latter possibility is supported by the *in silico* and biophysical observations made by Casilano and colleagues (*13*), identifying the 234 glycan as essential for Spike function. Since N234A Spike protein and BOA still interact *in vitro*, as seen by our SEC results, glycans other than the one attached at 234 must play a role in the interaction.

What picture emerges from our observations? Our *in vitro* data suggest that BOA binds sugars on multiple Spike proteins, possibly crosslinking different Spike proteins on a single virion or crosslinking Spike proteins on different virions (Figure 6). Such a mechanism is compatible with results obtained for the 2G12 antibody, which binds to glycans at positions 709, 717, and 801 of the Spike protein but fails to inhibit viral infection (*43, 44*). Because 2G12 is a monovalent glycan-binding antibody (*44*), it is not able to crosslink the Spike protein, in contrast to BOA and OAA that can bind in a multi-site mode. Therefore, we posit that BOA inhibition of SARS-CoV-2 infection is associated with simultaneous binding to multiple Spike proteins. This mechanistic model is further supported by our finding that a monovalent BOA variant binds the Spike protein but does not form soluble aggregates *in vitro* and fails to inhibit viral entry.

**Figure 6:**
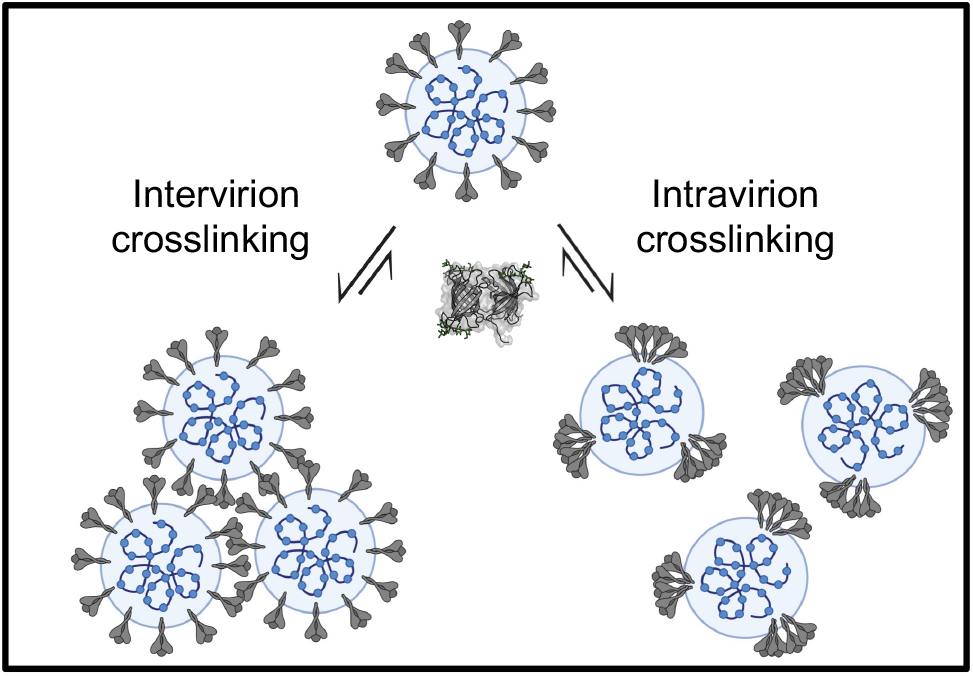
Model of BOA-mediated crosslinking of Spike proteins on SARS-CoV-2 (created with BioRender.com)

SARS-CoV-2 variants of concern possess minimal changes with respect glycosylation (*32*). For this reason and because BOA inhibition of viral entry is mediated through interactions with glycans on the Spike protein, BOA is a potent inhibitor against variants of concern. Our finding that BOA also inhibits an evolutionarily similar virus, SARS-CoV 2003, demonstrates that the inhibitory activity of BOA is minimally dependent on protein sequence. Since BOA inhibits viral entry of all tested SARS-CoV-2 variants, as well as SARS-CoV 2003, our findings may open the way to develop and exploit multivalent glycan binders in the quest to generate pan-coronavirus inhibitors.

## Conclusions

The potent inhibitory activity of the BOA lectin towards SARS-CoV-2 was demonstrated and characterized. This activity persists against variants of concern and SARS-CoV 2003, illustrating the tantalizing potential to develop multi-site glycan binders as pan-coronavirus inhibitors.

## Materials and Methods

### Protein expression and purification

#### BOA

*E. coli* BL21(DE3) cells were transformed with a pET15b plasmid harboring the gene for BOA; the protein was expressed with an N-terminal hexa-His tag and purified as previously described (*24*). In short, 100 mL of Luria-Bertani (LB) broth containing100 μg/ml of carbenicillin was inoculated with a single colony and incubated overnight at 37°C. 10 mL of the overnight culture was added to 0.5 L of LB containing 100 μg/ml carbenicillin, followed by incubation at 37°C until an optical density at 600 nm of 0.6 was achieved. Protein expression was then induced with 1mM IPTG, followed by incubation at 18°C for 16 hrs. Cells were harvested by centrifugation at 3,000 × g, and the cell pellet was resuspended in 50 mL of 20 mM NaH_2_PO_4_, pH 8.0, 300 mM NaCl, 10 mM imidazole, 1 mM TCEP. The cells were lysed using a microfluidizer, and cell debris was removed by centrifugation for 45 minutes at 18,000 × g. The supernatant was loaded onto a Cytiva HisTrap HP column and eluted using an imidazole gradient from 0-100% (20 mM NaH_2_PO_4_ pH 8.0, 300 mM NaCl, 500 mM imidazole, and 1 mM TCEP). Fractions containing BOA were dialyzed overnight at 4°C against 50 mM Tris-HCL pH 8.0, 375 μM EDTA, 1 mM TCEP, and the His-tag was cleaved by overnight incubation with tobacco etch virus protease (1:20 TEV to BOA ratio) and removed by passage over a HisTrap HP column. The flow-through containing BOA was concentrated to ~2 mL and loaded onto a S75 size-exclusion column and separated in 20 mM NaPO_4_ pH 7.5. Fractions containing BOA were collected, concentrated and frozen at −20°C. Purity and identity of the BOA protein was assessed by SDS-PAGE and LC-MS.

#### OAA

E. coli BL21DE3 cells were transformed with a pET26b plasmid containing the gene for OAA, and the protein was expressed following the same procedure as was used for BOA. OAA was purified as previously described (*22*). Briefly, cells were pelleted at 3000 × g and resuspended in 20 mM Tris-HCl pH 8.5, lysed using a microfluidizer, and centrifuged at 18,000 × *g* for 45 minutes to remove cellular debris. This lysate was dialyzed overnight into 20 mM Tris-HCl pH 8.5 (at 25°C) and loaded onto a Q HP anion exchange column and eluted using a 0-1 M gradient of NaCl + 20 mM Tris-HCl. Fractions were analyzed via SDS-PAGE and fractions containing OAA were concentrated using an Amicon spin concentrator. OAA was further purified by SEC chromatography using 20 mM Tris-HCL, 100 mM NaCl pH 8.5 (at 25°C). Fractions containing OAA were collected, concentrated and frozen at −20°C until further use.

#### Spike protein

The pαH vector harboring the gene for Hexapro S was obtained from Addgene (Plasmid #166856) and used to transfect expi293F cells as previously described (*30*). Cells were induced for expression with 3.5 mM valproic acid, and media containing the secreted Spike protein was collected after 3 days and clarified by centrifugation. Clarified supernatant was passed over a HisTrap HP column and eluted by a 0-100% gradient of elution buffer (20 mM NaH_2_PO_4_ pH 8.0, 150 mM NaCl, 500 mM imidazole). Fractions were analyzed for the presence of Spike protein Spike on SDS-PAGE, and Spike protein-containing fractions were pooled and concentrated to < 1mL. Further purification was carried out by size exclusion chromatography over a Superose-6 10/300 GL column in 20 mM NaPO_4_, 150 mM NaCl, pH 7.5, and Spike protein-containing fractions were pooled, concentrated, and flash frozen in 100 μL aliquots. Purity and identity of the Spike protein was confirmed by SDS-PAGE. Negative stain electron microscopy was used to confirm the Spike protein was in the pre-fusion conformation.

### Analytical size exclusion chromatography

100μL samples of purified Spike protein (1 μM) in the absence or presence of 5 or 10 μM BOA were passed over a Superose-6 10/300 GL column in 20 mM NaPO_4_, 150 mM NaCl, pH 7.5, using a flowrate of 0.8 mL/min over one column volume. Data were exported and plotted in MatLab.

### Dynamic light scattering

Light scattering data were acquired at 298 K using a DynaPro Plate Reader III (Wyatt Technology, Santa Barbara, CA) in a 384-well plate format. Samples (60 μL total volume in 20 mM NaPO_4_, 150 mM NaCl, pH 7.5) contained 2 μM Spike protein and increasing BOA concentrations (32 nM to 2 μM). For each sample ten sets of scattering data were collected and averaged in Dynamics 7.1.7. Data for each concentration were collected in triplicate. Hydrodynamic radii were extracted, and the data was analyzed in MatLab. The errors are given as the standard deviation of the mean for 3 replicates.

### Negative stain electron microscopy

Carbon-coated grids (Electron Microscopy Sciences, Hatfield, PA) were glow discharged for 30 seconds at 10 mA using the PELCO system (Ted Pella, USA). Grids were prepared by applying a 2 μl protein sample to the glow-discharged grid, followed by removal of excess liquid after 30 sec, and staining with 2% Uranyl acetate. The grids were allowed to dry for 15 minutes before use. All negative stain EM images were obtained on a Tecnai TF20 (FEI Company) equipped with a TVIPS CMOS camera.

### Fluorescence binding experiments

BOA was incubated with 2 × molar excess of Setau-405-maleimide (Seta biomedicals; Cat# K7-548) in 20 mM NaPO_4_, 150 mM NaCl or 12-16 hours. The reaction was quenched by filtering through a 10 kDa cut-off spin concentrator, with three buffer exchanges into fresh 20 mM NaPO_4_, 150 mM NaCl). Labeling was confirmed by LC-MS. Binding was followed by incubating 125 nM BOA with increasing amounts of Spike protein (31, 62, 125, 250, 500, and 1000 nM). Fluorescence intensity was normalized to the maximum fluorescence. Errors are given as the standard deviation of the mean for triplicate measurements. Since no 1:1 or defined stoichiometry of binding exists, the data was not fit to a model and only the half point of the saturation was extracted.

### NMR Spectroscopy

All NMR spectra were collected at 37°C using a Bruker AVANCE 600 MHz Spectrometer equipped with a triple resonance cryoprobe. 2D ^1^H-^15^N HSQC spectra were collected on 250 μM samples of BOAd3WRE1 in 20 mM NaAcetate pH 5.0 with 10% D_2_O. Glycan binding was assessed by addition of 250 μM 3α,6α-mannopentaose. All spectra were processed in NMRFAM-Sparky (*45*).

### Generation of Wuhan-Hu-1 pseudoviruses

Replication-defective pseudotyped HIV-1 encoding firefly luciferase was produced by transfection of 293T cells with plasmids encoding the SARS-CoV-2 Spike (Wuhan-Hu-1 with a 21 amino acid C-terminal deletion and the D614G substitution (HDM-SARS2-S-del21-D614G) (*46*) and the HIV-1 provirus (NLdE-luc) (*47*) as previously described. The N165A and N234A variants of HDM-SARS2-S-del21-D614G were generated by site-directed PCR mutagenesis using the QuickChange II kit (Agilent), and changes were confirmed by sequencing. Specific infectivity, the luciferase actvity normalized to the amount of virus, was determined by measuring the luciferase activity of virus stocks pseudotyped with WT, N165A, or N234A Spike in Vero E6 cells and quantifying HIV-1 capsid (p24) concentrations by ELISA (XpressBio).

Pseudotyped replication-defective lentivirus encoding a GFP reporter gene and expressing D614G SARS-CoV-2 Spike protein or the corresponding Spike proteins of the Alpha, Beta, Gamma, Delta, and Omicron variants of concern were purchased from Integral Molecular (catalog #s RVP-702, RVP-706, RVP-724, RVP-708, RVP-763 and RVP-768G).

### Pseudovirus inhibition assays

Infections of Vero E6 cells with pseudoviruses encoding luciferase were performed in triplicate for each condition in two independent experiments in 96-well plates. Eight dilutions of BOA, or no BOA (100% infectivity), were used, as previously described (*48*). For GFP encoding pseudovirus infection assays, HEK293F cells expressing a clonally stable hACE2 were obtained from Integral Molecular (cat# c-HA102) and cultured with puromycin to induce hACE2 expression. Infection assays were performed in 96-well plates with BOA concentrations ranging from 15 nM to 2 μM, generated by serial dilution with Dulbecco’s Modified Eagle Media (DMEM). Each well contained 50 μL of BOA in DMEM, to which 20 μL of virus and 30 μL of DMEM were added; these along with infection control wells (20 μL of virus to which 80 μL of DMEM was added) were incubated at 37°C for 1 hour. HEK293F cells were cultured to 80% confluency and lifted from the plate by trypsinization for 3 minutes, followed by addition of 20 mL DMEM, centrifugation at 90 × g for 10 minutes. The cell pellet was resuspended in fresh DMEM. Cell concentrations were determined using nucleocounter NC-100, and stock solutions of 2 × 10^5^ cells per mL were prepared. To each virus-containing well, 100 μL of cell stock solution was added. Plates were incubated for 72 hours at 37°C to allow for infection and expression of GFP. After 72 hours, the supernatant in each well was removed by gentle aspiration, and 200 μL of PBS were added. The plate was subsequently imaged using a bottom reading plate reader with 488 nm excitation and 535 nm emission filters. All variants were imaged in triplicate, and the imaging data were exported to MatLab. The fluorescence signal was normalized to control cells without BOA and fit to a sigmoidal function. The error bars represent standard deviations from the mean of triplicate or duplicate experiments.

### SARS-CoV-2 live-virus neutralization assays

BOA was diluted to 240, 200, 160, 120, 80, 40, 20, 16, 12, 8 or 4 nM in Opti-MEM. 100 μL of BOA at the different concentrations was mixed with 100 μl of SARS-CoV-2 inoculum (50 plaque forming units of virus in Opti-MEM) to yield final concentrations of 120, 100, 80, 60, 40, 20, 10, 8, 6, 4, 2 nM BOA. These mixes were used in inhibition assays as previously described (*49*) in Vero E6 cells for the Munich isolate or Vero-hACE2-TMPRSS2 cells for Delta and Omicron variants.

### SARS-CoV 2003 live-virus neutralization assays

Full-length SARS-CoV 2003 urbani expressing nanoluciferase (nLuc) was used as described previously (*50, 51*). One day prior to the assay, Vero E6 USAMRID cells were plated at 20,000 cells per well in clear-bottom black-walled plates. BOA was tested at a starting concentration of 2μg/ml and was serially diluted in equal volume of diluted virus and incubated at 37 °C with 5% CO_2_ for 1 h. After incubation, BOA and viral complexes were added at 800 plaque-forming units (PFU) at 37 °C with 5% CO_2_. Twenty-four hours later, plates were read by measuring luciferase activity using a Nano-Glo Luciferase Assay System (Promega) kit. Luminescence was measured by a Spectramax M3 plate reader (Molecular Devices). Virus neutralization titers were defined as the sample dilution at which a 50% reduction in relative light units was observed relative to the average of the virus control wells.

## Acknowledgements

The authors thank Teresa Bronsenitsch for her critical reading of this manuscript, Mike Delk for NMR support, and Kanako Mori for PCR mutagenesis to produce Spike mutants.

HDM_SARS2_Spike_del21_D614G was a gift from Jesse Bloom (Addgene plasmid # 158762; http://n2t.net/addgene:158762; RRID:Addgene_158762). SARS-CoV-2 S HexaPro was a gift from Jason McLellan (Addgene plasmid # 154754; http://n2t.net/addgene:154754; RRID:Addgene_154754).

## Funding

This work was supported by an NIH grant P50AI150481 (A.M.G) and the University of Pittsburgh Clinical and Translational Science Institute and the DSF Charitable Foundation (ZA). A.J.G was a Merck Fellow of the Life Science Research Foundation. RSB was supported by U54 CA260543. D.R.M is funded by a Hanna H. Gray Fellowship from the Howard Hughes Medical Institute. A.J.G. and D.R.M hold Postdoctoral Enrichment Program Award from the Burroughs Wellcome Fund.

All data needed to evaluate the conclusions in the paper are present in the paper or supplementary information. All data and reagents are available upon request from the authors.

