## Supplemental Information for "Targeting Spike Glycans to Inhibit SARS-CoV2 Viral Entry"

Table S1: BOA $EC_{50}$ values in SARS-CoV-2 and pseudovirus neutralization assays.

| **Virus** | **$EC_{50}$** |
|---|---|
| SARS-CoV-2 Munich isolate | 9 ± 1.0 nM (plaque assay) |
| SARS-CoV-2 Delta | 14 ± 0.5 nM (plaque assay) |
| SARS-CoV-2 Omicron | 11 ± 2.0 nM (plaque assay) |
| SARS-CoV 2003 Urbani | 26 ± 0.3 nM (Luciferase reporter assay) |
| Pseudovirus Wuhan-Hu-1 D614G S | 1.6 ± 0.3 nM (Luciferase reporter assay) |
| Pseudovirus D614G S | 48 ± 16 nM (GFP reporter assay) |
| Pseudovirus Alpha S | 76 ± 14 nM (GFP reporter assay) |
| Pseudovirus Beta S | 74 ± 21 nM (GFP reporter assay) |
| Pseudovirus Gamma S | 52 ± 7 nM (GFP reporter assay) |
| Pseudovirus Delta S | 43 ± 6 nM (GFP reporter assay) |
| Pseudovirus Omicron S | 70 ± 10 nM (GFP reporter assay) |

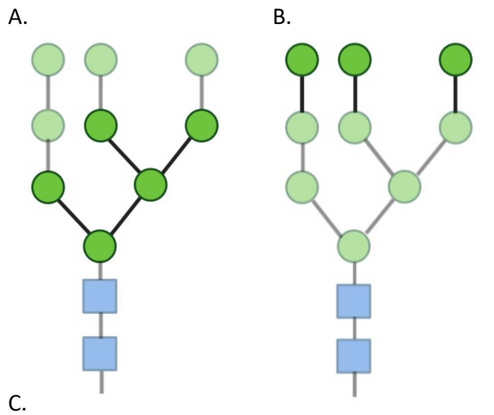


C.

| **BOA** | **OAA** | **GRFT** | **UB** |
|---|---|---|---|
| 115 nM | 75 nM | 80 nM | 5000 nM |
| 100% inhibition | Some inhibition | 0% inhibition | 0% inhibition |

Figure S1: A, B. Binding epitopes of BOA and OAA on (A) high mannose glycans and (B) GRFT. The interacting units are depicted by the dark green circles. C. Inhibition of SARS-CoV-2 infectivity by BOA, OAA, and GRFT. At ~100 nM, BOA was 100% inhibitory, while all others exhibited some or no inhibitory activity. Assay concentrations were chosen as four fold dilution from lectin concentrations where cytotoxicity was observed  Recombinantly expressed ubiquitin (UB) was used as a control.

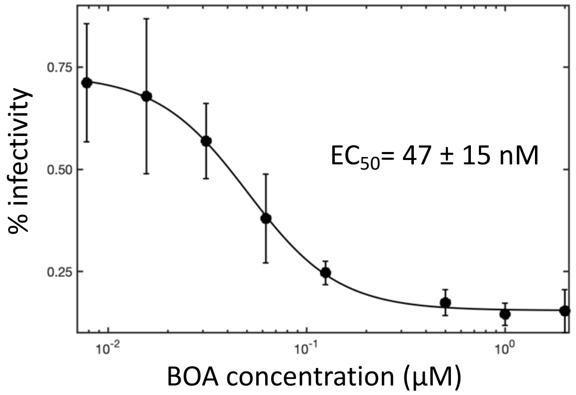


Figure S2: Viral inhibition using GFP expressing HIV-1 pseudotyped with D614G ancestral S, yielding an $EC_{50}$ value of 47 ± 15 nM.

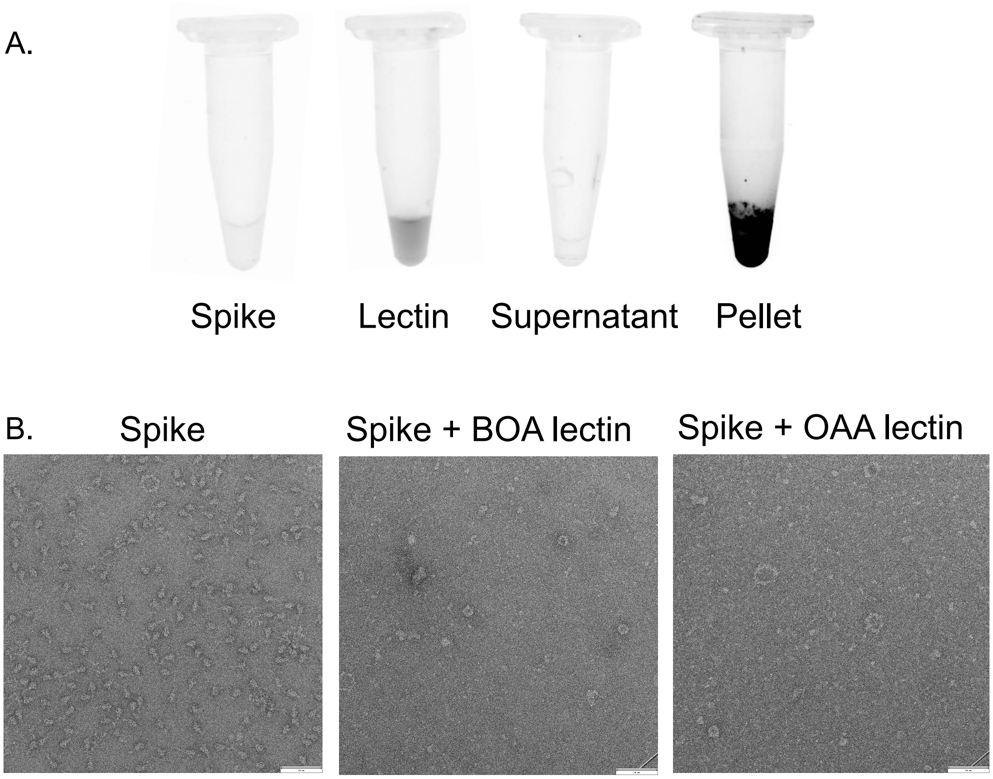


Figure S3: A. Precipitation assay of Spike protein with SeTau-405 labeled BOA. All fluorescent BOA accumulated in the precipitated complex in the pellet. B. Negative stain micrographs of Spike protein, Spike protein with BOA and Spike protein with OAA. Scale Bar, 70 nm

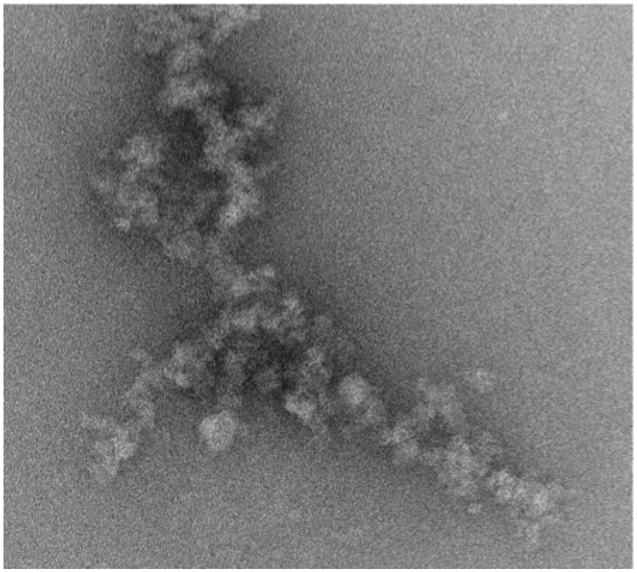

Figure S4: Negative stain micrographs of the soluble aggregates formed by the interaction between Spike protein and BOA, eluted from the Superose-6 10/300 GL.

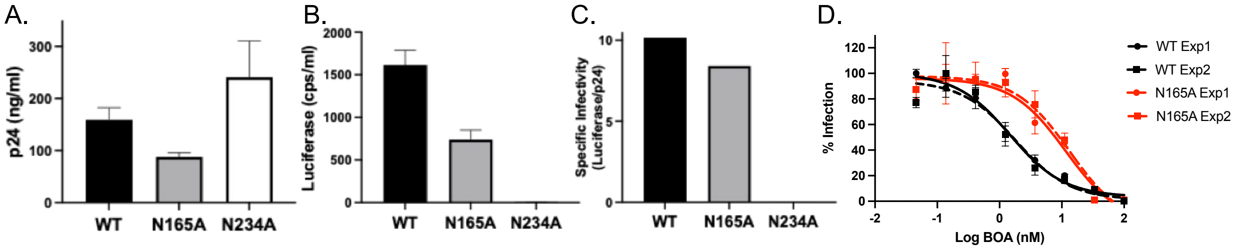


Figure S5: Characterization of HIV-1 pseudotyped with wild-type (WT) Spike (Wuhan-Hu-1) or N165A or N234A mutant Spike. (A) HIV-1 capsid (p24) production for all pseudotyped viruses confirmed virion production for all three variants. (B) Luciferase activity after infection of Vero E6 cells is shown. (C) Specific infectivity (infectivity normalized to virus production). Only viruses pseudotyped with WT and N165A showed activity, demonstrating that the N234A glycan deletion variant is noninfectious. (D) Inhibition of WT and N165A pseudoviruses by BOA is shown. The results show the averages and standard deviations of two independent experiments.

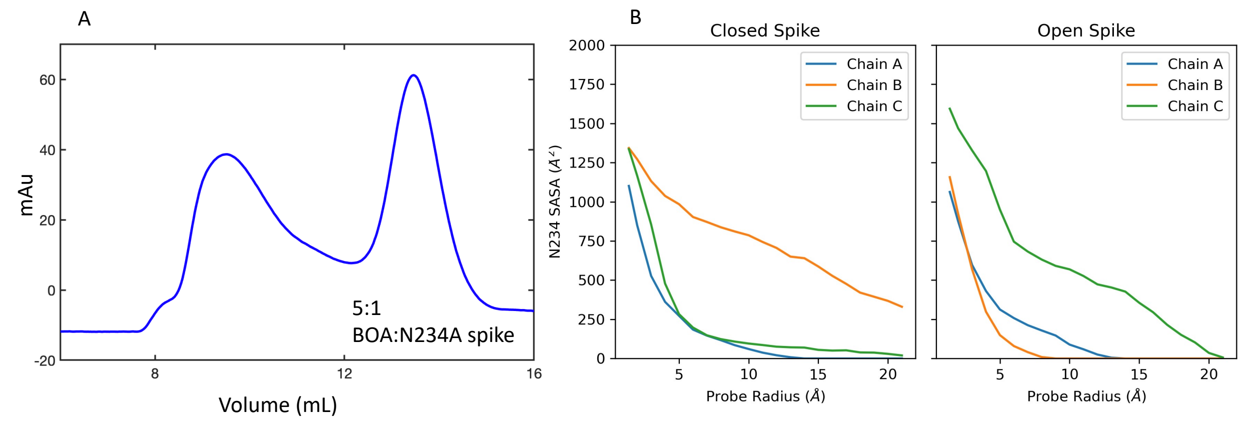


Figure S6: A. Size exclusion chromatogram of 1 µM N234A Spike protein in the presence of 5 µM BOA. A shift in elution profile indicates the formation of soluble aggregates. B. Solvent accessible surface area of the glycosylated Spike protein structure probed with spheres of radii ranging from 1.4 Å to 20 Å. The size dependence suggests that the N234 glycan is buried in each Spike protein chain in the trimeric structure and therefore inaccessible to molecules with radii larger than 20 Å, such as BOA.

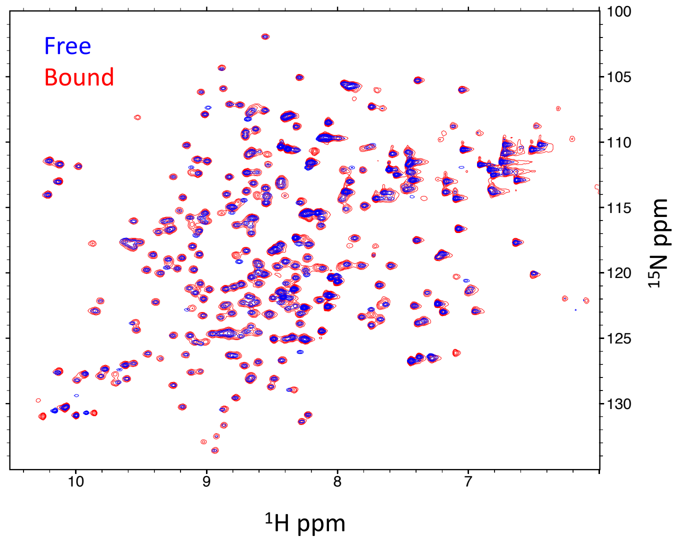


Figure S7: Superposition of the $^{1}$H,$^{15}$N HSQC NMR spectra of the BOAd3WRE1 variant in the absence (blue) and presence (red) of the Man5 core sugars (3α,6α-mannopentaose). The well-dispersed resonances, similar to the WT resonances, confirm that the overall protein structure is not changed and that the protein can bind glycan via its single remaining binding site.
